# Leveraging machine learning and citizen science data to describe flowering phenology across South Africa

**DOI:** 10.1101/2023.12.21.572952

**Authors:** R. D. Stewart, N. Bard, M. van der Bank, T. J. Davies

**Affiliations:** African Centre for DNA Barcoding (ACDB), Department of Botany and Plant Biotechnology, University of Johannesburg, P.O. Box 524, Auckland Park 2006, South Africa; Department of Biological & Agricultural Sciences, Sol Plaatje University, Kimberley, South Africa; Biodiversity Research Centre, University of British Columbia, Vancouver, British Columbia, Canada

**Keywords:** citizen science, flowering, iNaturalist, machine learning, phenology, phenological patterns

## Abstract

- Phenology — the timing of recurring life history events—is strongly linked to climate. Shifts in phenology have important implications for trophic interactions, ecosystem functioning and community ecology. However, data on plant phenology can be time consuming to collect and current records are biased across space and taxonomy.
- Here, we explore the performance of Convolutional Neural Networks (CNN) for classifying flowering phenology on a very large and taxonomically diverse dataset of citizen science images. We analyse >1.8 million iNaturalist records for plants listed in the National Botanical Gardens within South Africa, a country famed for its floristic diversity (∼21,000 species) but poorly represented in phenological databases.
- We were able to correctly classify images with >90% accuracy. Using metadata associated with each image, we then reconstructed the timing of peak flower production and length of the flowering season for the 6,986 species with >5 iNaturalist records.
- Our analysis illustrates how machine learning tools can leverage the vast wealth of citizen science biodiversity data to describe large-scale phenological dynamics. We suggest such approaches may be particularly valuable where data on plant phenology is currently lacking.

## Introduction

Phenological patterns, describing the temporal sequencing of recurring life history events, such as flowering, provide information on plant life cycles and reproductive strategies, species ecological interactions, and responses to shifts in the global climate (Forrest & Miller-Rushing, 2010; Richardson *et al*., 2013; Wolkovich *et al*., 2014b). It is now well recognised that many plant species are shifting their phenology in response to recent climate warming, flowering and fruiting earlier, and senescing later (Parmesan & Yohe, 2003; Menzel *et al*., 2006; Cleland *et al*., 2007; Thackeray *et al*., 2016). There is concern that such shifts might give rise to ‘phenological mismatches’ among interacting organisms (Miller-Rushing *et al*., 2010), with potential ecosystem consequences (Memmott *et al*., 2007; Duchenne *et al*., 2020). For example, temporal mismatches between flowering plants and their pollinators could result in reduced pollination success and seed set (e.g. Kudo and Ida 2013), leading to population declines (see Willis et al., 2008). However, to date, empirical evidence for species asynchrony is mixed (Kharouba *et al*., 2018) and constrained by data availability (Kharouba & Wolkovich, 2020).

While there is growing recognition that phenological responses can vary importantly across species (Wolkovich *et al*., 2012), data on plant phenology is often both taxonomically and geographical limited (Cook *et al*., 2012). The majority of phenological research is currently concentrated in the temperate regions of North America (Parmesan, 2007; Miller-rushing *et al*., 2011), Europe (Menzel *et al*., 2006; Duchenne *et al*., 2020), and Asia (Kudo *et al*., 2004; Ge *et al*., 2015), with only few studies in the southern hemisphere (e.g. Central and South America; Niu *et al*., 2013, Australia and New Zealand; Chambers *et al*., 2013; Rawal *et al*., 2015). Data from Africa are even more sparse (Adole *et al*., 2016; for exceptions, see Dreyer *et al*., 2006; Williams *et al*., 2021). To generate accurate ecological forecasts of climate change impacts on biodiversity, there is a critical need to rapidly scale-up data collection to fill this gap, and new approaches are needed.

Herbaria provide one valuable, non-traditional, source of phenological data (Miller-Rushing *et al*., 2006; Willis *et al*., 2017a; Meineke *et al*., 2018, 2019; Lang *et al*., 2019). The increasing digitization of herbarium and natural history collections records has facilitated access to historical datasets that document phenological shifts across time (see e.g. Primack *et al*., 2004; Soltis, 2017; Nelson and Ellis, 2019; Hedrick *et al*., 2020). However, this new data stream has given rise to new challenges: how to accurately document and record species phenology from sometimes many thousands of digitized images (e.g. Park *et al*., 2019; Lee *et al*., 2022). One approach has been to leverage citizen scientists to review and score digital images, such as implemented through the *CrowdCurio* crowdsourcing tool (Willis *et al*., 2017b). New Machine Learning (ML) techniques provide an alternative approach and when trained on well curated data have been shown to perform with high accuracy (Lorieul *et al*., 2019; Davis *et al*., 2020; Goëau *et al*., 2020). For example, Losrieul *et al*. (2019) utilized Convolutional Neural Networks (CNN) to classify images of herbarium specimens into specific phenophases, and Davis *et al*. (2020) used Region-based Convolutional Neural Networks (R-CNN) to detect and count the three reproductive stages of budding, flowering, and fruiting for six wildflower species. This latter study showed how a model trained on one species could be applied to another, albeit phylogenetically closely related, species. Pearson *et al*. (2020) explore and discuss some of the strengths and constraints inherent within such approaches. The potential of herbarium data to characterise the flowering phenology of plants in southern Africa has recently been demonstrated by Williams *et al*. (2021) and Daru *et al*. (2019), although neither explore the application of ML tools.

While herbaria have provided an opportunity to rapidly increase the taxonomic breadth of species for which we can obtain phenological data, the geographical origins of the specimens they contain remains highly skewed. Specimens tend to be collected close to roads and herbaria, and a few collectors (along with their individual collecting habitats) tend to dominate collections (Daru *et al*., 2018). Public databases populated by citizen scientist contributors present an additional source of ecological data that can be used by researchers (Dickinson *et al*., 2012). iNaturalist (Unger *et al*., 2021), provides a platform for contributors to submit observation data electronically to a centralized database. At the time of writing, the iNaturalist database contained 164,567,244 observations for 430,848 species, of which 44% were plants. Such databases also contain biases towards some regions and taxonomic groups, over-representing well-visited locations and more charismatic taxa (Di Cecco *et al*., 2021; Daru & Rodriguez, 2023). However, when analysed appropriately, the vast wealth of ecological data in such databases, including precise metadata on locations and times of observations, offers valuable insights into both temporal and spatial diversity patterns (Dickinson *et al*., 2010; Mesaglio *et al*., 2021).

iNaturalist images of plants capture their phenology, and Reeb *et al*. (2022) demonstrated the application of CNN to classify the phenology of garlic mustard (*Alliaria petiolate*) into distinct phenophases: vegetative, budding, flowering, and fruiting. In contrast to herbarium records, which are typically mounted on standardized herbarium sheets and digitized using fixed mounted cameras (see Gutiérrez-Larruscain *et al*., 2018), iNaturalist images are frequently of plants in their natural habitats, and capture considerable diversity in background environment, proximity to the camera, image resolution, and lighting (Barve *et al*., 2020). It might, therefore, be assumed that training ML tools on iNaturalist images would require an unreasonably large training dataset (Willi *et al*., 2019); however, Reeb *et al*. (2022) showed that their model algorithm accuracy was no different to manual classification. The application of ML tools to large citizen science databases, such as iNaturalist, thus has potential to massively scale up phenological records (Barve *et al*., 2020). Here, we explore this potential, applying CNN to classify flowering phenology of the plants of South Africa using images from iNaturalist.

South Africa is one of the world’s most biodiverse countries, with more than 21,000 plant species (Zengeya *et al*., 2020), encompassing three globally recognized biodiversity hotspots: the Cape Floristic Region, Succulent Karoo and Maputaland-Pondoland-Albany (Myers *et al*., 2000), and nine different biomes spanning various climate regimes, from the arid Karoo to the cool, wet climate of the Drakensberg. We target our analysis on the over 8,600 species recorded within and around National Botanical Gardens, where iNaturalist sampling is regionally richest. Our focus on National Botanical Gardens not only ensures a high density of images in iNaturalist, leveraging the known bias in sampling (Di Cecco *et al*., 2021), but includes sites within each biodiversity hotspot and four of the nine biomes (Grassland, Savanna, Fynbos, and Albany ticket).

Previous studies have demonstrated high accuracy of ML algorithms in scoring flowering phenology using well curated datasets, often for a single species, and using standardised images. Our analysis evaluates the performance of CNN in classifying flowering phenology across a large number of species with highly diverse flowering morphology from citizen science photographs frequently taken in natural settings. We validate model accuracy by comparing our phenological classifications to independent data on expert reported phenology from Manning & Goldblatt (2012).

## Methodology

We explored the performance of convolutional neural networks (CNNs) in classifying flowering phenology from citizen science images on the iNaturalist digital platform. In contrast to the narrow taxonomic focus of most previous studies, we tackle a diverse array of plant species from South Africa, one of the world’s most biodiverse floras.

We centre our study on species occurring within (or with ranges projected to overlap with) the 11 South African National Botanical Gardens (NBG) as the plants within these gardens receive many visits from the public, and thus tend to be well represented on iNaturalist. We compiled species list from a variety of different sources including species distribution models (Hoveka *et al*., 2020), iNaturalist observation records (downloaded from the-national-botanical-gardens-of-southern-africa project; https://www.inaturalist.org/ on 21/03/2023), plants with historical collections in the areas (Magill *et al*., 1983; Mucina *et al*., 2000; Germishuizen & Meyer, 2003; Rutherford *et al*., 2003, 2012; Henderson, 2011), and a list provided by Botanic Gardens Conservation International (BGCI; https://www.bgci.org/). For purposes here, we excluded species within the order Poales, which tend to have less conspicuous flowers and are less well-represented in iNaturalist, returning a final list of 11,272 species within 1,626 genera across 204 families.

We cleaned species names by removing author abbreviations, checked for spelling inconsistencies using The Plant List (http://www.theplantlist.org/), and making sure that the final names agreed with accepted names obtained from Plant of the World Online (https://powo.science.kew.org/). We next downloaded from iNaturalist (GBIF DarwinCore Archive; https://www.inaturalist.org/pages/developers; 25/07/2023) all research grade observation records in South Africa for this taxon set, providing us with a total of 1,807,310 images. For observation to be classified as research grade, there must be at least a 2/3 consensus on the identification of the species pictured.

Initial testing of the efficiency of the classification model highlighted that charismatic fruits were occasionally misclassified as flowers, thus we implemented a two-step process to classify flowering phenology: first, a primary model that classified images as not flowering vs flowering/fruiting, and a secondary model to differentiate between flowering and fruiting. The secondary model was only run on the images that were classified as flowering/fruiting in the primary model.

### Constructing the training dataset

The training datasets were constructed using a random sample of species from iNaturalist, including no more than two images per species per category to maximise taxonomic representation. Images were manually sorted into three categories: (1) not flowering or fruiting, (2) fruiting, and (3) flowering. The primary training set consisted of two categories (flowering and non-flowering) each consisting of 5,000 images. The secondary training dataset was constructed from a subset of the primary flowering/fruiting dataset and consisted of 1,000 images of which 168 were fruiting. The overall accuracy of the classification pipeline was evaluated on an independent dataset of 2,000 pre-classified images (1,000 flowering, 500 fruiting, and 500 not flowering or fruiting).

### Training the Convolutional Neural Networks – ResNet-18

A CNN is a specialized model well-suited for processing visual data. It uses layers that detect features in images, each layer tuned to specific image features such as edges, textures, or shapes through convolutional filters. ResNet-18 is a type of deep convolutional neural network known for its 72-layer architecture with 18 deep layers, featuring residual blocks with skip connections (Chandola *et al*., 2021). It is commonly used in computer vision tasks due to its ability to efficiently train deep models while maintaining high performance. We use ResNet-18 because it has fewer convolutional layers, making it less computationally intensive compared to other pre-trained neural networks. ResNet-18 underwent pre-training on the ImageNet dataset, consisting of 1,281,167 training images (Deng *et al*., 2010).

Before classification, we resized iNaturalist images to an input of 300 × 300 pixels and normalized the distribution of RGB values to a range between zero and one. Each image was associated with the metadata provided by iNaturalist including date, family, longitude, and latitude (Table S1).

Both models were trained on their respective training datasets, with 80% of the data used for training and the remaining 20% of data used for model validation. We optimised the learning rate (0.01, 0.001, 0.0001 and 0.00001), dropout rate (0.4, 0.3, 0.2 and 0.1), and n dense (1024, 512, 256 and 128) separately for each model using the training dataset. We then compared models varying number of Epochs (20, 50 100 or 200) and batch size (16, 32 and 64), evaluating model performance using the evaluation dataset (2,000 pre-classified images).

The final primary model used the following variables: epochs = 50, batch size = 32, learning rate = 0.001, drop outrate = 0.4 and n dense = 1024. The final secondary model used the following variables: epochs = 25, batch size = 32, learning rate = 0.001, drop outrate = 0.4 and n dense = 512.

### Model classification of iNaturalist images

We first employed the primary model to classify each of the 1,807,310 images as flowering/fruiting or not flowering/fruiting. Images with flowering/fruiting prediction probability values >0.5 were then run through the secondary model to distinguish between flowering and fruiting.

To describe the local phenology of the 11 National Botanical Gardens, we selected iNaturalist records using a rectangular bounding box around each garden (Table S2). We then generated separate density plots of the flowering periods for each garden.

### Evaluating the accuracy and precision of the modelled phenology

The accuracy of the two-step method was evaluated by cross referencing our phenological classifications with independent expert-reported phenology data from Manning & Goldblatt (2012). For species matching in both datasets, we recorded: (1) the number of images classified as flowering that fell within the flowering phenology window reported in Manning & Goldblatt – “true positives”; (2) the number of images classified as flowering that fell outside the Manning & Goldblatt phenological window – “false positives”; (3) the number of images classified as not flowering that fell inside the Manning & Goldblatt phenological window – “false negatives”; (4) the number of images classified as not flowering that fell outside the Manning & Goldblatt phenological window – “true negatives”. Importantly, although we use standard terminology to describe congruence between data sources, disagreement with Manning & Goldblatt (2012) does not necessarily indicate error in image classification by our CNN. We thus manually checked a selection of the images identified as “false positives” and “false negatives” to determine if these were correctly classified by our CNN model or represented true classification errors.

### Computational Information

The bulk of the coding was scripted using the R programming language for statistical computing (v4.3.0) and the neural networks were trained using Tensorflow (v2.7) in Python (v3.10). We used R packages data.table (v1.14.8), dplyr (v1.1.2), tidyverse (v2.0.0), lubridate (v1.9), stringr (v1.5.0), Taxonstand (v2.4), and taxize (v0.9.100) for data cleaning and dataframe manipulation. We conducted statistical analyses using R packages circular (v0.4-95), NPCirc (v3.1.1), and Hmisc (v5.1-0). The Python functions were called within R using packages keras (v2.11.1), tensorflow (2.11.0) and reticulate (v1.28). GGplot2 (v3.4.2) was used for data visualisation. All code can be found at https://github.com/rossdstewart/ML-Phenology-Code.

Image classification took approximately 50 ms/image on an Intel (R) Xeon (R) CPU e-2420 v2 @ 2.2GHz with 24 cores and 47 GB of system memory. The bulk of the image classification was conducted over a period of 48 days in batches of ∼2,000 species to minimise loss in event of unforeseen disruptions in processing.

## Results

We analysed over 1.8 million images representing 11,272 species, of which 54.77% (989,874) were classified as flowering (Table S1). The best represented families were Asteraceae, Iridaceae Proteaceae and Fabaceae, with each of these families contributing >100,000 images. Of the top 20 families, Iridaceae had the largest percentage (83.66%) of flowering images (Table 1).

**Table 1.** Details of image classification for the top 20 best represented families in iNaturalist. Bottom entry provides the equivalent values summed across all (204) families (Table S3).

| **Family** | **Species** | **Species flowering** | **% species recorded flowering** | **Flowering images** | **Not flowering images** | **% images flowering** |
| --- | --- | --- | --- | --- | --- | --- |
| **Asteraceae** | 1,368 | 1,350 | 98.68 | 144,349 | 77,818 | 64.97 |
| **Iridaceae** | 821 | 817 | 99.51 | 94,320 | 18,427 | 83.66 |
| **Proteaceae** | 371 | 366 | 98.65 | 76,477 | 81,207 | 48.5 |
| **Fabaceae** | 1,037 | 1,002 | 96.62 | 71,419 | 87,842 | 44.84 |
| **Ericaceae** | 516 | 508 | 98.45 | 66,370 | 17,928 | 78.73 |
| **Orchidaceae** | 325 | 320 | 98.46 | 36,422 | 11,976 | 75.26 |
| **Aizoaceae** | 756 | 694 | 91.80 | 35,195 | 21,439 | 62.14 |
| **Geraniaceae** | 207 | 201 | 97.10 | 35,094 | 18,912 | 64.98 |
| **Oxalidaceae** | 119 | 116 | 97.48 | 27,225 | 9,061 | 75.03 |
| **Scrophulariaceae** | 337 | 333 | 98.81 | 24,438 | 10,711 | 69.53 |
| **Asparagaceae** | 399 | 379 | 94.99 | 23,299 | 24,020 | 49.24 |
| **Campanulaceae** | 169 | 168 | 99.41 | 22,500 | 6,705 | 77.04 |
| **Crassulaceae** | 229 | 203 | 88.65 | 21,208 | 25,240 | 45.66 |
| **Amaryllidaceae** | 211 | 204 | 96.68 | 18,219 | 12,414 | 59.48 |
| **Malvaceae** | 223 | 213 | 95.52 | 16,580 | 13,507 | 55.11 |
| **Lamiaceae** | 185 | 183 | 98.92 | 16,044 | 11,298 | 58.68 |
| **Thymelaeaceae** | 126 | 125 | 99.21 | 15,719 | 7,166 | 68.69 |
| **Apocynaceae** | 318 | 298 | 93.71 | 15,716 | 20,929 | 42.89 |
| **Asphodelaceae** | 317 | 286 | 90.22 | 13,252 | 20,565 | 39.19 |
| **Rutaceae** | 187 | 181 | 96.79 | 12,441 | 7,872 | 61.25 |
| **ALL (204) families** | **11,272** | **10,809** | **95.89** | **989,874** | **817,436** | **54.77** |

Our two-step method (primary plus secondary models) was able to correctly classify 90.8% of the 2000 pre-categorized images (Fig. 2a). While the primary model alone was able to correctly classify 90.2% of the pre-categorized images. A total of 6,195 taxa were shared with the expert-reported phenology data from Manning & Goldblatt (2012). From this shared taxon set, 570,958 (41.62%) images classified as flowering fell within the reported phenological window reported by Manning & Goldblatt (2012) – “true positives” (Fig. 2b), and of the 589,392 images classified as non-flowering 268,687 (45.59%) fell within the non-flowering period reported by Manning & Goldblatt (2012) – “true negatives”. Notably, however, 214,269 (27.22%) of the images classified as flowering fell outside of the reported phenological window for the species. Importantly, while we denote these mismatches as “false positives”, in many instances they represent true records of flowering (correctly classified images) that occur beyond the current records of flowering times (for examples see Fig. 3), and thus extend our knowledge of flowering phenology for many taxa.

**Fig. 1.**
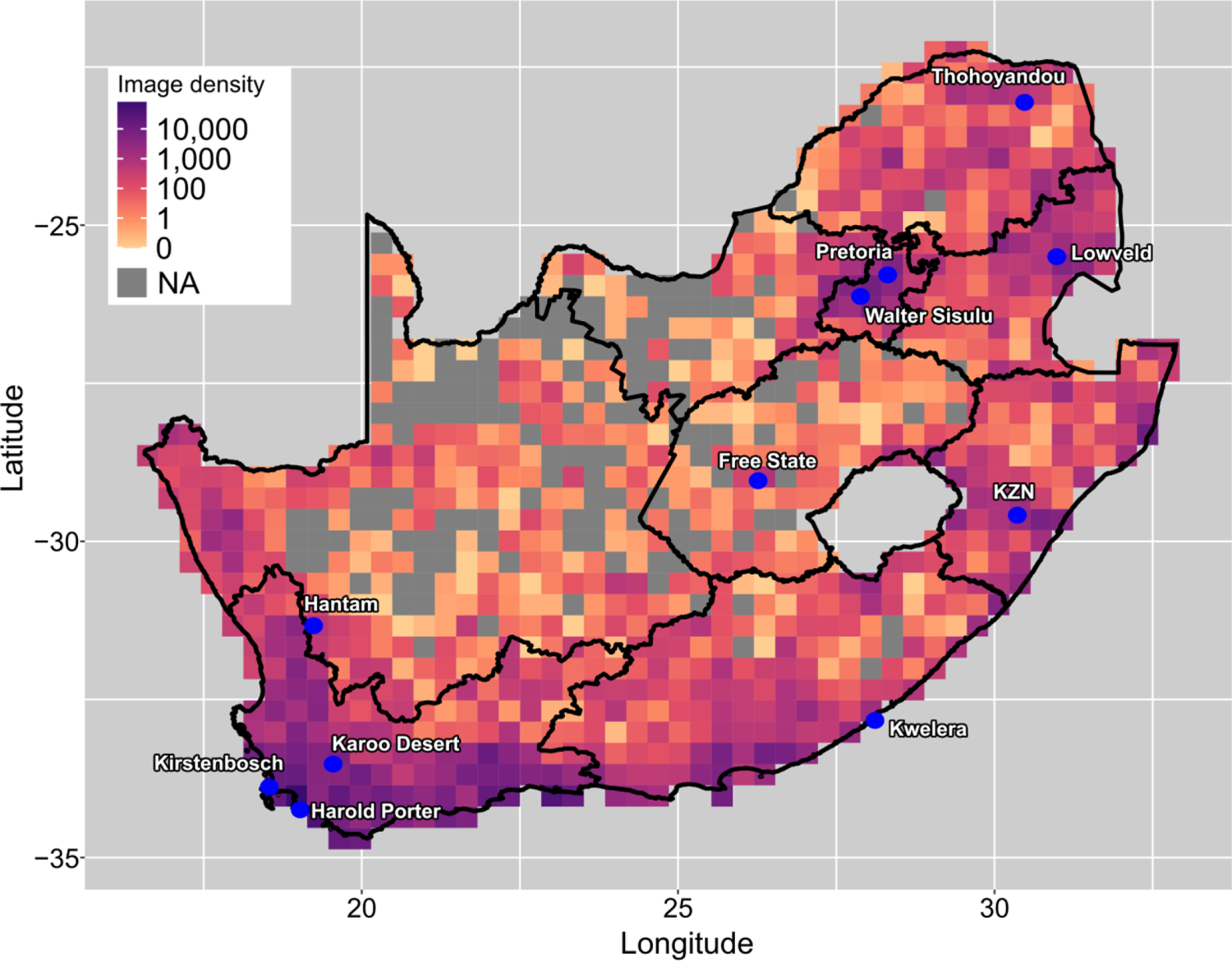
Density and geographical coverage of research grade iNaturalist images (n=1,807,310) across South Africa. Locations of the National Botanical Gardens in South Africa indicated with blue symbols, map shading proportional to the density of iNaturalist images within 0.33° grid cells (darker colours = higher density). (GBIF DarwinCore Archive from https://www.inaturalist.org/pages/developers; 25//07/2023)

**Fig. 2.**
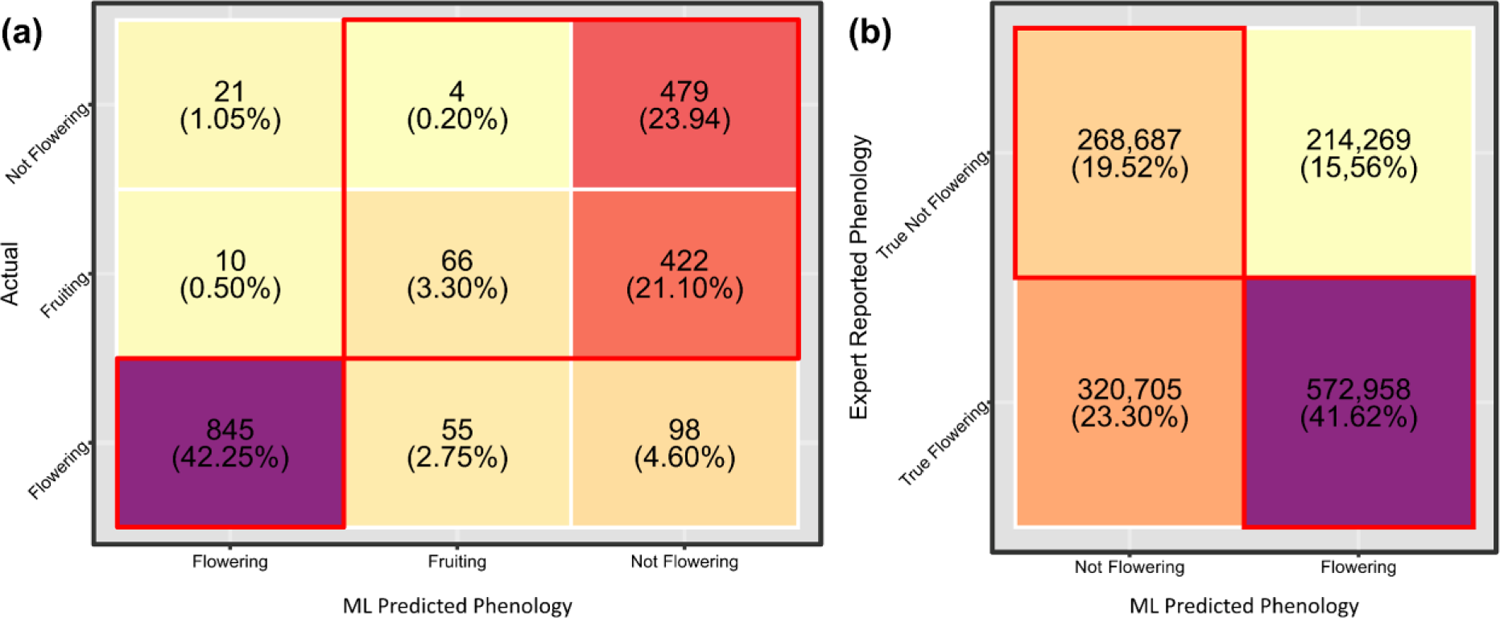
Confusion matrices for (a) the evaluation image dataset (n = 2,000) representing manually categorized images (not flowering, fruiting and flowering) versus CNN classified images using our two-step method; and (b) the phenological patterns predicted from this study vs independent expert-reported phenology from Manning & Goldblatt (2012). Shading indicates strength of agreement between the two datasests (darker colours = higher agreement). Values in cells indicate number of images.

**Fig. 3.**
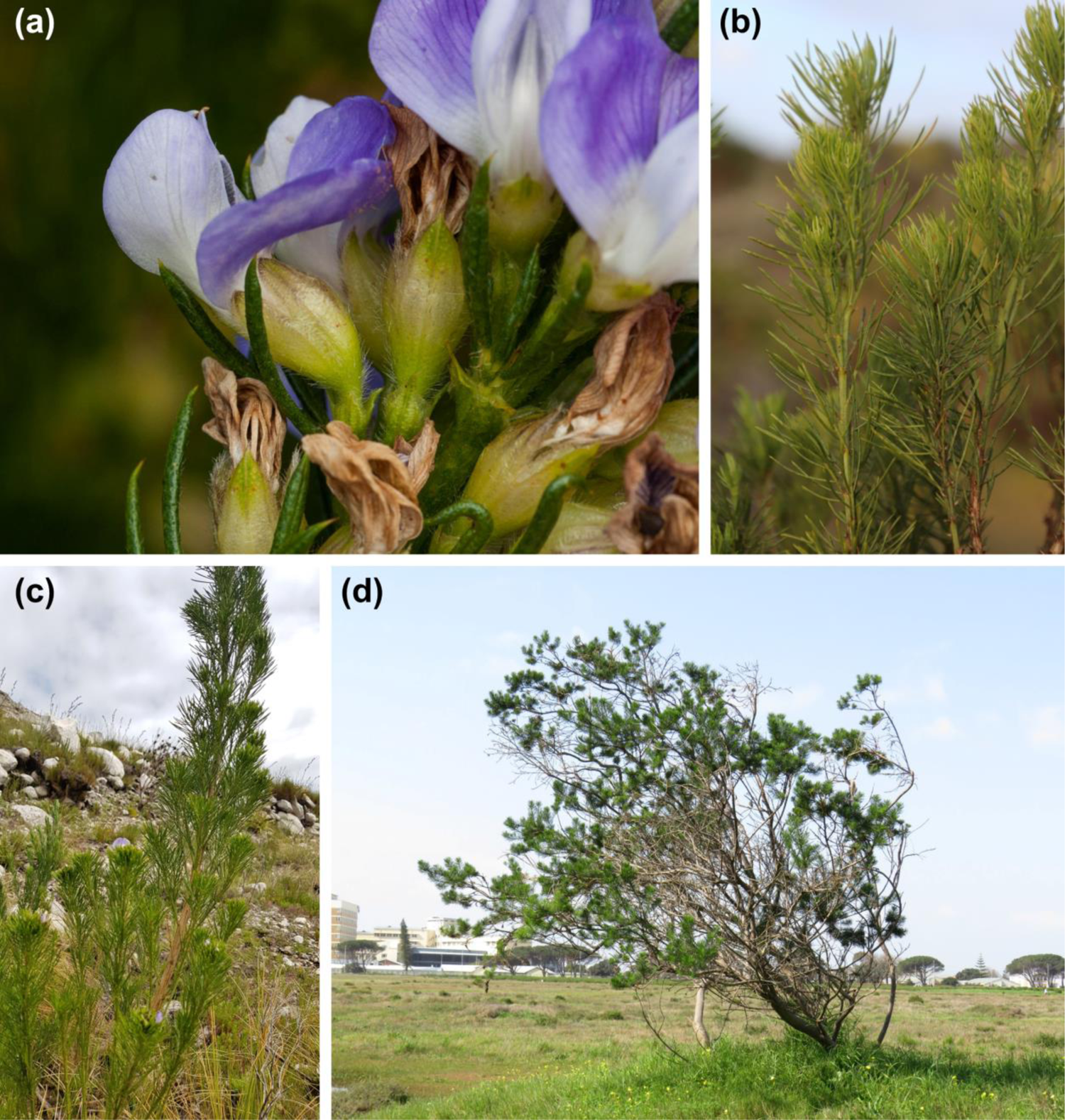
Examples images of *Psoralea pinnata* classified as flowering/non-flowering that were outside the expert-reported phenological window for the species. (a) image correctly catogrised as flowering (© magriet b; iNat photo ID: 15712292; 2013); (b) image incorrectly catogrised as flowering (© Tony Rebelo; iNat photo ID: 90138261; 2020); (c) image incorrectly catogrised as non-flowering (© suretha_dorse; iNat photo ID: 97811160; 2020) and (d) image correctly catogrised as non-flowering (© Jeremy Gilmore; iNat photo ID: 58287501; 2019).

We mapped the temporal distribution of flowering times by fitting a circular density function to the unique observations of flowering for each of the 6,986 species with >5 records of flowering (Fig. 4), a threshold that allowed us to reconstruct reasonable estimates of peak flowering times. Individual density plots were fit assuming a smoothing bandwidth (bw) = 50, kernel = “vonmises”, and removing duplicate observations. Across species, clusters in peak flowering can be observed at the end of April and then from the end of August though to the end of October (all image classifications found in Table S1). Geographically, we observe two distinct flowering phenologies: in the west the mean flowering time falls around September and in the east mean flowering centres around the end of April (Fig. 5a). However, there is larger variation in flowering phenology along coastal habitats, most noticeable towards the south of the country (Fig. 5b).

**Fig. 4.**
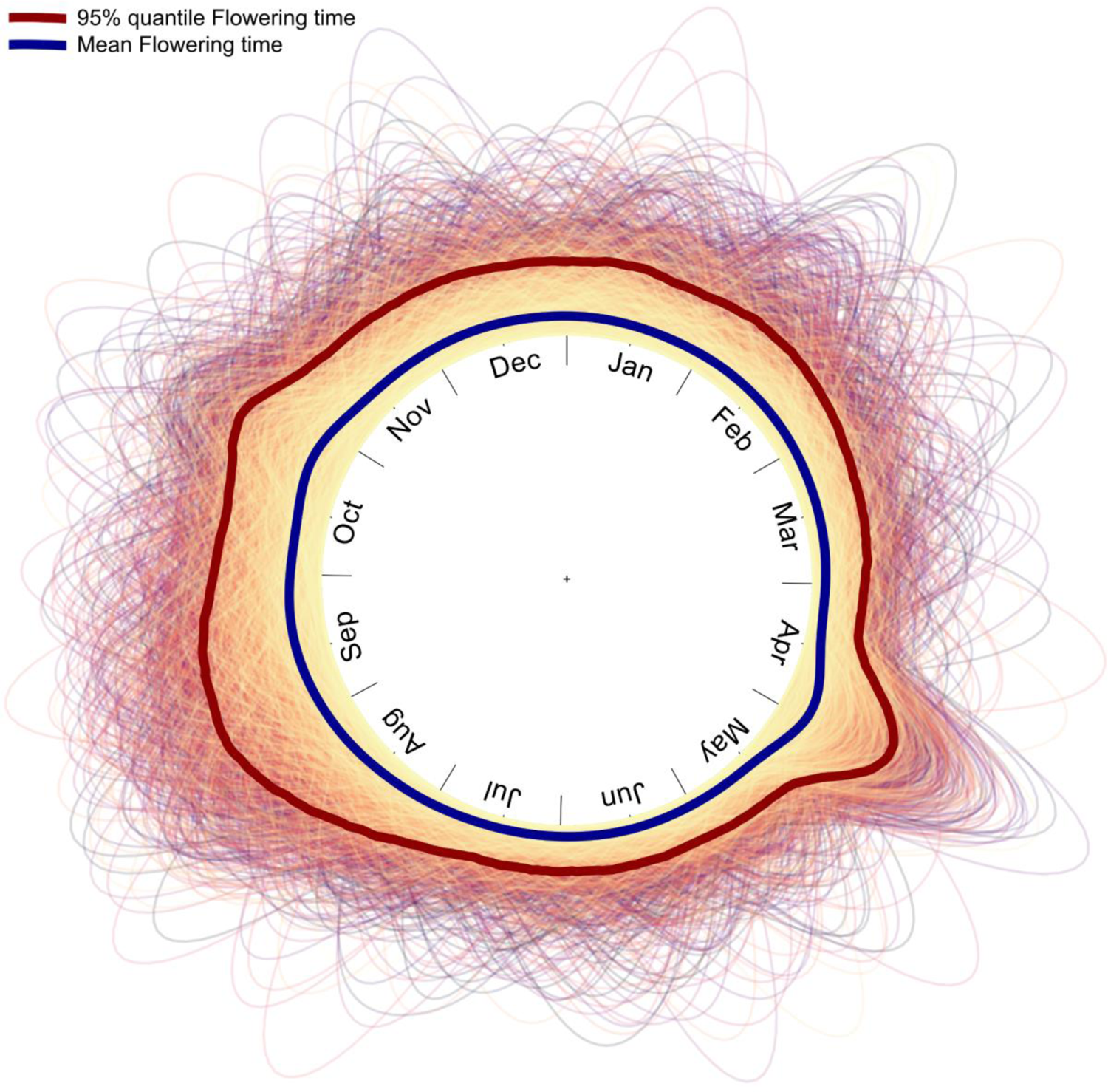
Circular density plots of flowering times for species with >5 images categorized as flowering (6,986 species, representing a total of 497,911 flowering images). The blue line is the mean, and the maroon line shows the 95% quantile. Equivalent plot for all evaluated species based on the classification of 1,807,310 images can be found in Table S1.

**Fig. 5.**
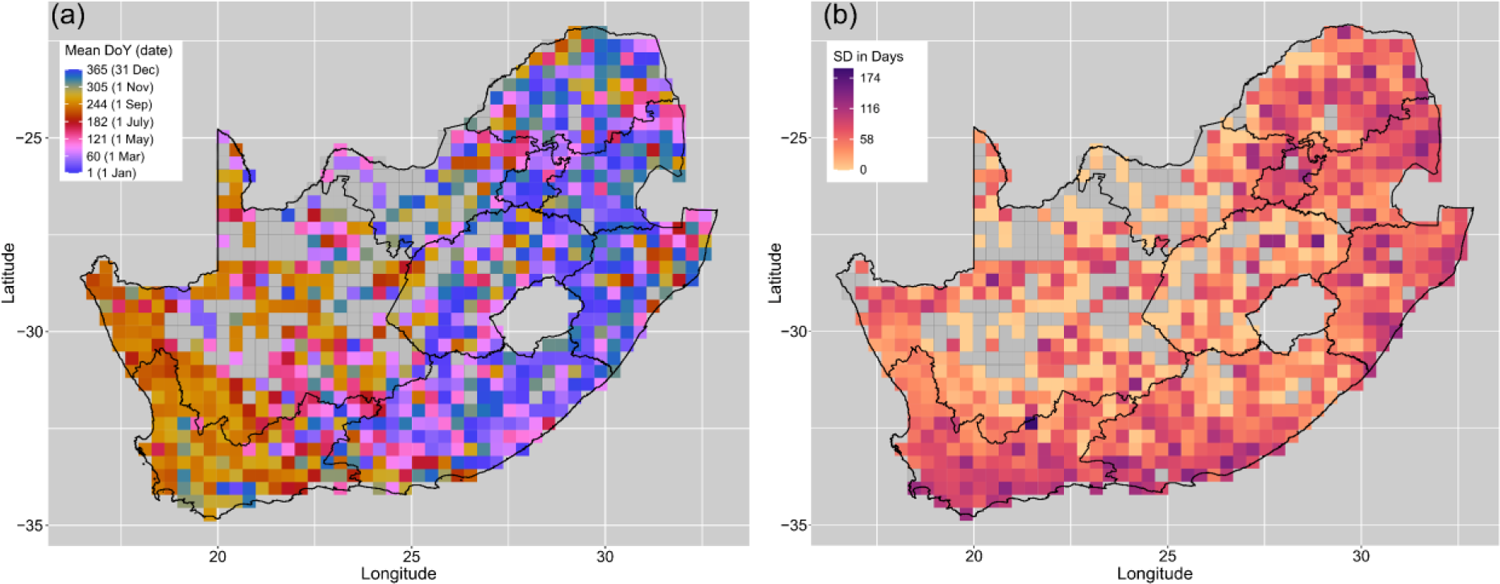
Flowering phenology of South African plants represented within National Botanical Gardens. (a) Mean day of year (DoY) of flowering. (b) Standard deviation (SD) of flowering, in days.

The geographical variation in flowering phenology is further emphasised in comparisons of peak flowering among National Botanical Gardens (Fig. 6). For example, in Lowveld NBG (images=81, species=45) the peak flowering period is during the winter months (May-July), while Thohoyandou (images=38; species=19), also in the savanna biome, has a pronounced peak earlier in the year at the end of April, although this is estimated from a lower sample size of images and species. Harold Porter NBG (images=2,026, species=312) and Kirstenbosch NBG (images=5,256; species=545), which are both located within the Fynbos biome, show broadly similar phenological patterns, but with the former showing a more distinct secondary peak in flowering later in the year.

**Fig. 6.**
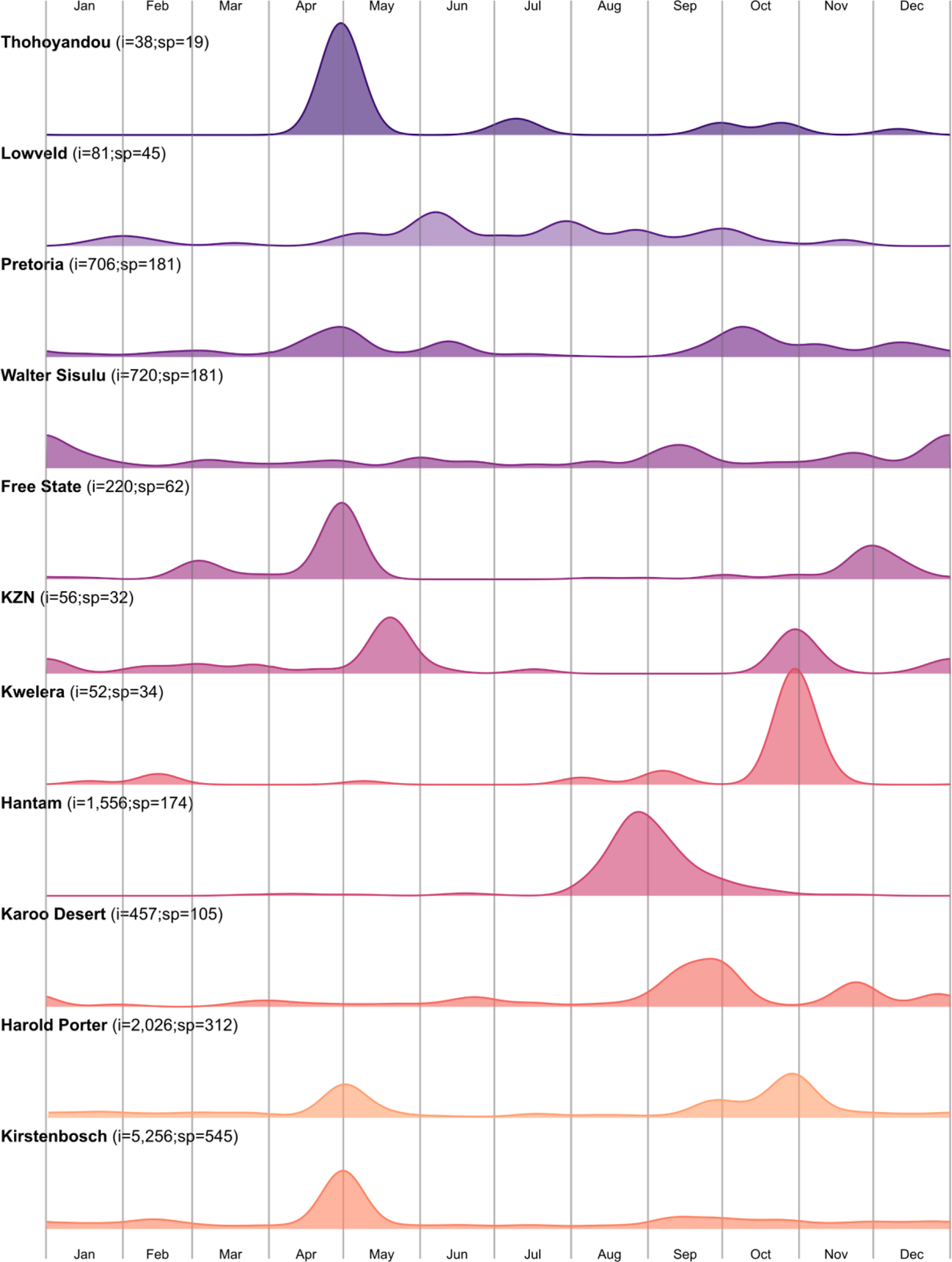
Local phenology of the 11 National Botanical Gardens in South Africa using the two-step method on the set of images from iNaturalist selected using a bounding box around each botanical garden (Table S2). i = number of images; sp = number of species.

## Discussion

We evaluated the performance of convolutional neural networks (CNN) in classifying flowering phenology from citizen science images on the iNaturalist digital platform. Previous work has demonstrated the potential of machine learning algorithms, such as CNNs, in extracting phenological data from digital images, but have generally applied methods to a single or a few closely related species using carefully curated datasets (e.g. Pearson *et al*., 2020; Soltis *et al*., 2020; Love *et al*., 2021; Davis *et al*., 2022). Here, we apply a CNN classifier (ResNet-18) to the flora of South Africa, a region famed for its exceptional floristic diversity. In contrast to most existing studies, we use a training dataset composed of one to two images from several thousand different species, capturing large variation in flowering morphology and plant architecture, from trees to herbs. Images also vary from landscape panoramas to macrophotographs of flowers. Our model was able to correctly classify images with over 90% accuracy, a similar performance to other recent phenological annotation models using more standard approaches (e.g. Reeb *et al*. 2022: 95.9%, Lorieul *et al*. 2019: 96.3%, and Pearson *et al*. 2020: 87.8%).

Using metadata associated with each image on iNaturalist, including day of year the image was recorded, we compared the temporal overlap between our phenological classifications to independent data on expert-reported phenology from Manning & Goldblatt (2012). Manning & Goldblatt (2012) report the flowering period (months) for over 7,157 species of the Greater Cape Floristic Region, providing one of the most comprehensive sources of flowering phenology for South Africa. Of the images we classified as flowering, 73% fell within the phenological window described by Manning & Goldblatt (2012) for that species, a much lower percentage than estimated when evaluating model accuracy using our evaluation dataset of 2,000 pre-classified images. Thus, over a quarter of images we classified as flowering occur outside of the expected phenological flowering period for the species; this does not, however, necessarily indicate a failure in our CNN classifier. A manual check of a random selection of images mismatched with Manning & Goldblatt (2012) verifies that many images were correctly classified.

Manning & Goldblatt (2012) report flowering times for species within the Greater Cape Floristic Region; however, many species from the Cape region have distributions outside the Cape. Because flowering phenology is largely a response to environmental cues, including sometimes complex interactions between temperature, rainfall and photoperiod (Flynn & Wolkovich, 2018; Wolkovich *et al*., 2022), we would expect the same species to flower at different times in different locations if those location also differ in cues (Schwartz *et al*., 2006). The iNaturalist images we examine were drawn from across South Africa, and it is likely that we expand the geographical sampling of some Cape species beyond the Cape region. It is perhaps no surprise, therefore, that we expand the temporal window of flowering phenology for these species as we increase the environmental niche space encompassing their distributions. Not only do these records add to our knowledge of flowering times, but they also provide information on the underlying cues driving flowering phenology. For example, by matching temperature and precipitation records to flowering observations, we can vastly expand current phenological models across taxa and environmental space (Wolkovich *et al*., 2013; Sandor *et al*., 2021).

It is also possible that some of the discrepancy between our flowering records and the phenology reported by Manning & Goldblatt (2012) captures the imprint of recent climate change. It has been more than a decade since Manning & Goldblatt (2012) published their flora for the Cape, and some species may have shifted in their phenologies since these data were originally compiled. For instance *Gladiolus monticola* G.J.Lewis ex Goldblatt & J.C.Manning, a narrow ranged endemic commonly found on Table Mountain, a popular tourist attraction, is recorded as flowering from mid-December to February, and occasionally in March (Goldblatt & Manning, 2020). However, we observe flowering from January to July, with a peak in April (1,981 iNaturalist flowering images), suggesting this species may have shifted its phenology to later in the year. Although such recent phenological shifts are likely to be generally small, iNaturalist data may provide both critical baseline data to track future phenological changes and information on present day phenologies that can be contrasted with historical observations. For instance, in a herbarium study, Williams *et al*. (2021) examined 129 *Pelargonium* species (6,200 herbarium specimen) between 1901 – 2009, and found that peak flowering had moved earlier by 11.6 days. While the temporal span of iNaturalist data is relatively short, over the last three years our data (images=19,346; mean species=139) indicates that peak flowering time has moved to later in the year (Fig. S1), highlighting large interannual variation in phenology that could complicate attempts to quantify historical trends.

While we have emphasised how our classifications importantly expand current knowledge of flowering phenology for species across South Africa, we also recognise that automated classifiers, such as CNNs, will almost always misclassify some images. In our model, just under 10% of images were misclassified. We used a flowering prediction threshold of 0.5, and errors of commission (classifying non-flowering images as flowering – false positives) could be lowered by increasing this threshold. For example, of the 6,195 species shared between our analysis of iNaturalist images and Manning & Goldblatt (2012) ∼3% had a flowering image categorization probability below 0.6. However, increasing the prediction threshold would obviously come at the cost of increasing errors of omission (failure to correctly classify flowering images as flowering – false negatives). There is, therefore, a trade-off between inflating false positives versus false negatives, and where to balance this trade-off will depend on the question of interest. When describing species phenological distributions from noisy data, we suggest one sensible approach would be to be to omit data points that lie outside a given standard deviation from the peak concentration of records as these are more likely to be erroneous, and we have provided code in the supplement for this purpose, using circular statistics. However, if constructing predictive models of phenology, it would also be possible to simply weight observations by our confidence in them, thus avoiding the need to threshold on predictions.

Our study is one of the first to provide a comprehensive database on flowering phenology for South African plants. Aggregating across species, we show evidence for separate peaks in flowering phenology around April (Autumn) and October (spring). We also show how mean flowering times vary spatially, highlighting the distinct flowering phenology in the western (winter rainfall) and eastern (summer rainfall) parts of the country. In the Cape Floristic Region, characterised by winter rainfall, spring marks the height of blooming, and there is a gradual decline in flowering towards autumn and early winter (Johnson, 1993). However, with the absence of freezing temperature that characterise temperate biomes, some species are still recorded flowering through winter months (Dreyer *et al*., 2006; Daru *et al*., 2019). In the summer rainfall, region species tend to flower through spring and summer (Venter & Witkowski, 2019; Fitchett & Raik, 2021). A number of studies in the winter rainfall region identify precipitation a key predictor of flowering phenology (Mayer & Kuhlmann, 2004; Dreyer *et al*., 2006; Wessels *et al*., 2011), while temperature may be more important in the summer rainfall savannas (Chidumayo, 2001). Both differences in timing of cues across space and regional variation in species sensitivities to alternative cues likely contribute to spatial variation in peak flowering times (Pau *et al*., 2011; Wolkovich *et al*., 2014a).

Examining the temporal sequence of flowering intensity across the National Botanical Gardens provides greater nuance. For example, some gardens are characterised by multiple peaks in flowering, such as KZN NBG and Free State NBG, indicating variation in sensitivity to climate cues across taxa. Other gardens, such as Lowveld NBG and Kirstenbosch NBG, have barely discernible peaks, with a low level of flowering through most of the year, perhaps reflecting deliberate planting to extend the flowering season for visitors to these gardens. We can also detect some evidence for a latitudinal gradient in flowering times with an earlier start to peak flowering at Hantam NBG in the North, and later peaks in flowering as we move southwards to the Karoo Desert NBG and then Harold Porter NBG. Thohoyandou, in Limpopo Province is one of South Africa’s youngest National Botanical Gardens, and has an unusually pronounced peak in flowering at the end of April; however, many of these species were found to be alien invasives, which pose a significant challenge to the province (Mbedzi *et al*., 2018; Moshobane *et al*., 2022).

Our analysis is both a demonstration of the potential for citizen science data to contribute to phenological research, especially when combined with machine learning, and a call for greater contributions to iNaturalist by citizen scientists. Currently, records are concentrated around more developed areas and contributors tend to focus disproportionately on some taxa (Di Cecco *et al*., 2021). In South Africa, we show that the geographical coverage of images is best in highly populated areas and in the Western Cape Province, while coverage remains sparse across much of the interior of the country. As iNaturalist and similar citizen scientist initiatives continue to grow, we hope these gaps will gradually shrink. With advances in machine learning tools it will also become possible to extract additional ecological information from images (Soltis *et al*., 2020), including data on plant functional traits (Younis *et al*., 2018; Weaver *et al*., 2020; Triki *et al*., 2022), community membership (Onishi & Ise, 2021), and plant health or disease status (Barbedo, 2019; Liu & Wang, 2021).

## Supporting information

Bounds of the gardens that were used to filter iNaturalist records to estimate local phenological patterns.

Details of image classification for all families.

Phenology of genus Pelargonium for the years 2019, 2020, and 2022 determined from iNaturalist images using the two-step method.

## Acknowledgements

We would like to thank Lerato Hoveka (species distribution models), Leslie Powrie (historical records) and Suzanne Sharrock (Botanic Gardens Conservation International) for assistance when complaining the lists of species from the National Botanical Gardens. We thank Paul Herbert for valuable feedback on an earlier version of this manuscript.

## Author contributions

R.D.S. conceptualized the method; R.D.S., M.vdB. and T.J.D. planned and designed the research questions; N.B. downloaded all images; R.D.S., N.B., and T.J.D. constructed and built the models. R.D.S. analysed the data; R.D.S. and T.J.D. interpreted results; R.D.S. and T.J.D. wrote the manuscript with significant input from N.B. and M.vdB.

## Data availability

All data and R code needed to recreate analyses are available on GitHub at https://github.com/rossdstewart/ML-Phenology-Code and at doi: xx

## Supporting Information

**Fig. S1** Phenology of genus *Pelargonium* for the years 2019, 2020, and 2022 determined from iNaturalist images using the two-step method.

**Table S1** Classification results of each image and associated meta data.

**Table S2** Bounds of the gardens that were used to filter iNaturalist records to estimate local phenological patterns.

**Table S3** Details of image classification for all families.

