## Supplementary figures and images for "Leveraging machine learning and citizen science data to describe flowering phenology across South Africa"

### Phenology of genus Pelargonium for the years 2019, 2020, and 2022 determined from iNaturalist images using the two-step method.

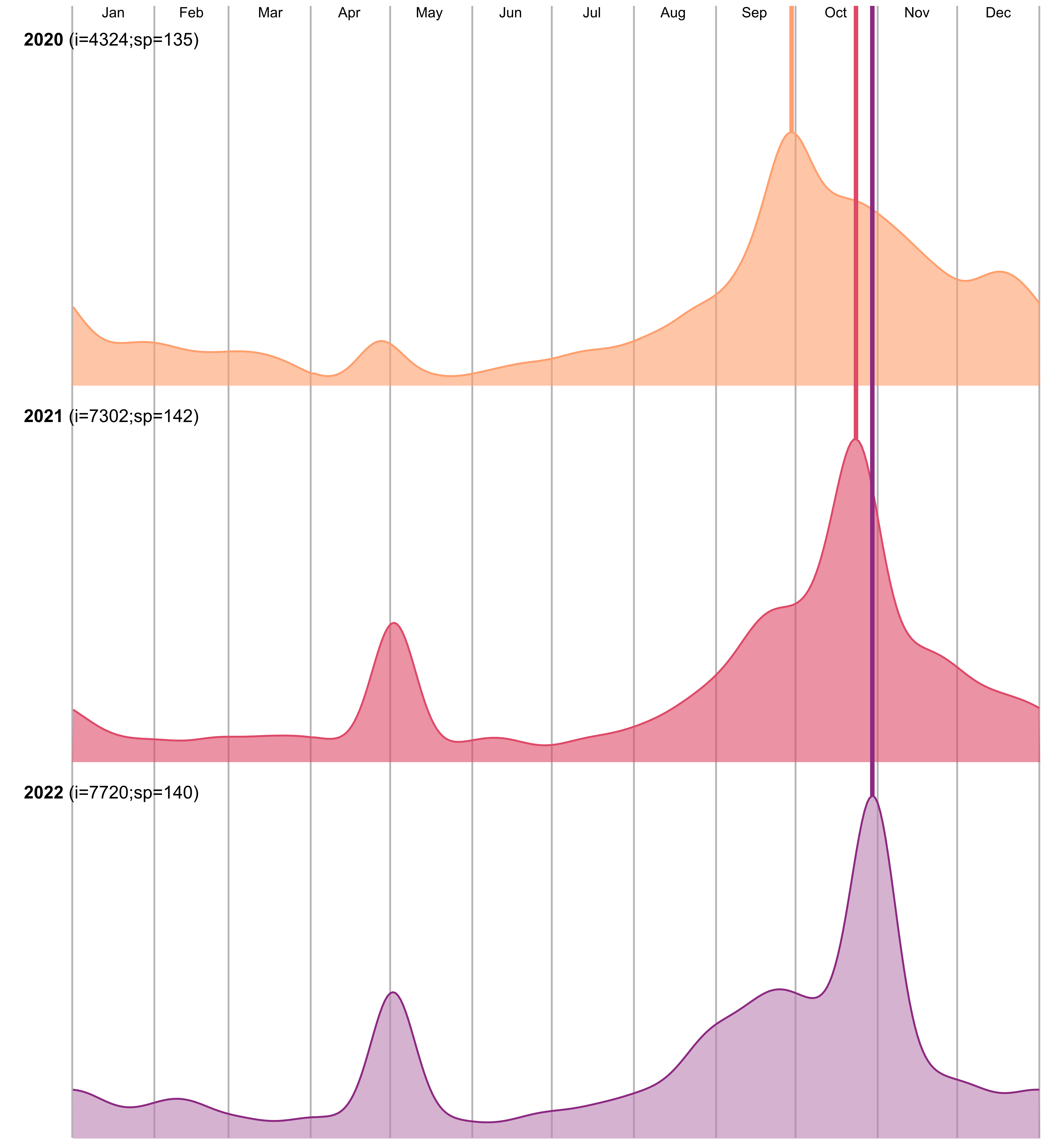
